# Analysis of genetic relatedness between gastric and oral *H. pylori* in patients with early gastric cancer using multilocus sequence typing

**DOI:** 10.1101/2022.08.08.503124

**Authors:** Ryoko Nagata, Hiroki Sato, Shoji Takenaka, Junji Yokoyama, Shuji Terai, Hitomi Mimuro, Yuichiro Noiri

## Abstract

Although oral cavity is the second most colonized site after the stomach, the association between gastric and oral *Helicobacter pylori* remains unclear. This study aimed to compare the genetic relatedness between gastric and oral *H. pylori* in Japanese patients with early gastric cancer by multilocus sequence typing (MLST) analysis using seven housekeeping genes. Gastric biopsy specimens and oral samples, including saliva, supragingival dental biofilm, and superficial layers of the tongue, were collected from 21 patients positive for *H. pylori* by a fecal antigen test. The number of *H. pylori* allelic profiles of seven loci obtained from oral and gastric samples ranged from zero to seven since the yield of DNA was small even when the nested PCR was performed. The alleles of seven loci from both collection sites were determined from only one patient, and two out of seven alleles matched between oral and gastric samples. MLST analysis revealed that only one sample had a matching oral and gastric *H. pylori* genotype, suggesting that different genotypes of *H. pylori* inhabit the oral cavity and gastric mucosa. The phylogenetic analysis showed that oral *H. pylori* in two patients was markedly similar to gastric *H. pylori*, implying that the the origins of two strains may be the same, and the stomach and the oral cavity may be infected at the same time. In brief, although different genotypes of *H. pylori* exist in the oral cavity independently of *H. pylori* present in the stomach, there are rare cases in which the same *H. pylori* is present in the stomach and the oral cavity. It is necessary to establish a culture method for oral *H. pylori* for elucidating whether the oral cavity will act as the source of the gastric infection, as our analysis was based on a limited number of allele sequences.

**Author summary:** *Hericobacter pylori* is a clinically important pathogen that causes chronic gastritis and peptic ulcers, which are associated with gastric carcinoma. Oral *H. pylori* DNA has also been detected in a range of oral specimens, including saliva, supra- or sub-gingival biofilm, dentin caries, dental pulp, infected root canal, and coating of the tongue. Thus, researchers speculate that the main route of transmission of gastric *H. pylori* infections is via the oral cavity, however, there is inadequate evidence to support this suggestion. This study aimed to investigate if the oral and gastric *H. pylori* have the same origin using multilocus sequence typing. The results of this study provided some insight into the role of oral cavity as the source of *H. pylori* infections in stomach.

## Introduction

*Helicobacter pylori* is a major causative pathogen of gastritis, peptic ulcers, and gastric carcinoma [1–4]. Transmission of *H. pylori* is thought to occur mainly via the oral cavity during childhood [5, 6]. *H. pylori* persists throughout a person’s life without specific therapy and most people infected with *H. pylori* usually remain asymptomatic. However, 30% of individuals may develop mild to severe upper gastrointestinal disease [6]. The prevalence of *H. pylori* infections among Japanese children and adolescents has been markedly decreasing with socioeconomical development. A meta-regression analysis of *H. pylori* infections in Japan from 1908–2003 revealed that the predicted prevalence in persons born in 1990 and 2000 was only 15.6% and 6.6%, respectively [7]. Thus, it is thought that the incidence of gastric cancer will continue to decrease over time [8].

As the oral cavity is the inlet port, researchers considered that it may also be the secondary habitat of *H. pylori* following colonization of the stomach. *H. pylori* grows spirally in the stomach, which is a microaerobic environment. In contrast, it is thought that it does not increase the number under aerobic or anaerobic conditions and they are viable but non-culturable in human oral cavity [13]. Therefore, nested polymerase chain reaction (PCR) is typically used to confirm their presence in oral specimens [9–12]. The specific DNA of *H. pylori* has been detected in a broad range of oral specimens, including saliva [12,14–17], supra- or sub-gingival biofilm [12,14,15,17,18], dentin caries [18], dental pulp [17, 19], infected root canal [18], and the coating of the tongue [12, 14]. We have reported that one-fifth of Japanese young adults have oral *H. pylori* DNA [12].

However, it has not been fully investigated whether *H. pylori* exists alive in the oral cavity and serves as a supply source for the stomach. The speculation that the oral cavity could also be a potential reservoir for *H. pylori* is based on epidemiological studies demonstrating a correlation between the presence of oral and gastric *H. pylori* [18,20,21]. However, there has been no evidence of a genotypic correlation between strains in the gastric mucosa and the oral cavity. Moreover, although there were some studies demonstrating that *H. pylori* in the oral cavity affected the outcome of eradication therapy [10], and that adjunctive periodontal therapy could enhance the efficiency of *H. pylori* treatment [22, 23], there is still no evidence that the recurrence after eradication therapy of gastric *H. pylori* is caused by oral *H. pylori*. In their systematic review and meta-analysis, López-Valverde et al. reported that there is no clear evidence that *H. pylori* present in oral bacterial biofilm causes gastric infections and vice versa [24].

Since there are few studies comparing genotype differences in multiple genes, we aimed to investigate the genetic relatedness between gastric and oral *H. pylori* in patients with *H. pylori* infection in the stomach. To overcome the difficulty of the analysis, we applied a multilocus sequence typing (MLST) technique. MLST, which consists of a combination of partial nucleotide sequences of several housekeeping genes, is valid to address the intraspecific phylogenetic structure between different strains of the bacteria [25, 26].

## Results

### Characteristics of patients

The characteristics of the participants are summarized in Table 1. We recruited 21patients (seven women and 14 men; mean age: 72.8 years, range: 59 to 86 years). Of the 21 participants, 20 had the adenocarcinoma in the stomach, and the other had the squamous cell carcinoma in the esophagus. The number of teeth and decayed, missing, and filled teeth (DMFT) ranged from 5–30 and 5–28, respectively. The average number of oral cleanings per day was 1.8, and 47.6% (10/21) in the patients used oral cleaning aids such as a dental floss and interdental brush. Further, 19.0% (4/21) of the participants were smokers.

**Table 1.** Demographic and clinical characteristics of the patients.

| Patient No. | Sex | Age | No. of remaining teeth | DMFT | Frequency of oral cleaning | Cleaning aid | Smoker | GI cancer |  |  |
| --- | --- | --- | --- | --- | --- | --- | --- | --- | --- | --- |
|  |  |  |  |  |  |  |  | Location | Histological type | Differentiation |
| 1 | F | 78 | 18 | 18 | 1 | - | - | Stomach | Adenocarcinoma | tub1>tub2 |
| 2 | F | 66 | 28 | 11 | 3 | + | - | Stomach | Adenocarcinoma | tub1, tub2 |
| 3 | M | 66 | 15 | 22 | 2 | - | - | Stomach | Adenocarcinoma | tub1 |
| 4 | M | 76 | 24 | 18 | 3 | - | - | Stomach | Adenocarcinoma | tub1 |
| 5 | F | 67 | 8 | 28 | 2 | + | - | Stomach | Adenocarcinoma | tub1 |
| 6 | F | 75 | 29 | 24 | 2 | + | - | Stomach | Adenocarcinoma | tub1 |
| 7 | M | 81 | 13 | 24 | 1 | + | - | Stomach | Adenocarcinoma | tub1>tub2 |
| 8 | M | 63 | 28 | 20 | 2 | - | + | Stomach | Adenocarcinoma | tub1 |
| 9 | M | 86 | 5 | 28 | 1 | - | - | Stomach | Adenocarcinoma | tub1 |
| 10 | M | 59 | 26 | 22 | 1 | - | + | Stomach | Adenocarcinoma | tub1 |
| 11 | M | 78 | 29 | 5 | 1 | - | - | Stomach | Adenocarcinoma | tub1>tub2 |
| 12 | F | 77 | 24 | 13 | 2 | + | - | Stomach | Adenocarcinoma | tub1, tub2 |
| 13 | M | 81 | 26 | 9 | 2 | + | - | Stomach | Adenocarcinoma | tub1 |
| 14 | F | 84 | 28 | 11 | 2 | + | - | Stomach | Adenocarcinoma | tub1 |
| 15 | F | 70 | 25 | 17 | 2 | + | - | Stomach | Adenocarcinoma | Por2, sig |
| 16 | M | 66 | 30 | 9 | 2 | + | + | Stomach | Adenocarcinoma | tub1, tub2 |
| 17 | M | 76 | 14 | 21 | 1 | - | - | Stomach | Adenocarcinoma | tub1>tub2 |
| 18 | M | 79 | 15 | 27 | 0 | - | - | Stomach | Adenocarcinoma | tub1 |
| 19 | M | 65 | 28 | 10 | 3 | + | - | Stomach | Adenocarcinoma | tub1, pap-<br>tub1>tub2 |
| 20 | M | 70 | 12 | 25 | 2 | - | - | Esophagus | Squamous cell carcinoma | Mod |
| 21 | M | 66 | 6 | 26 | 3 | - | + | Stomach | Adenocarcinoma | tub1, low grade |
| Mean± |  | 72.8 (± | 20.5 (± | 18.5 | 1.8 (±0.8) |  |  |  |  |  |
| SD |  | 7.6) | 8.4) | (±7.2) |  |  |  |  |  |  |

GI: gastrointestinal; Tub1: Tubular adenocarcinoma well differentiated type; Tub2: Tubular
adenocarcinoma moderately differentiated type; Por2: poorly differentiated adenocarcinoma non-solid type; Sig: Signet-ring cell carcinoma; Mod: Moderately differentiated; DMFT: Decayed, missing, and filled teeth.

### Histopathological observation of gastric tissue

A representative histopathological section was shown in Figure 1. Hematoxylin and eosin (H-E), and and Giemsa staining of the tissues showed spiral rods that penetrated between the gastric mucosa and mucus-producing cells. This is consistent with what has been reported as a characteristic of *H. pylori* [27, 28]. Bacteria with similar characteristics were also obtained from the culture of the stomach tissue of the patients.

**Figure 1.**
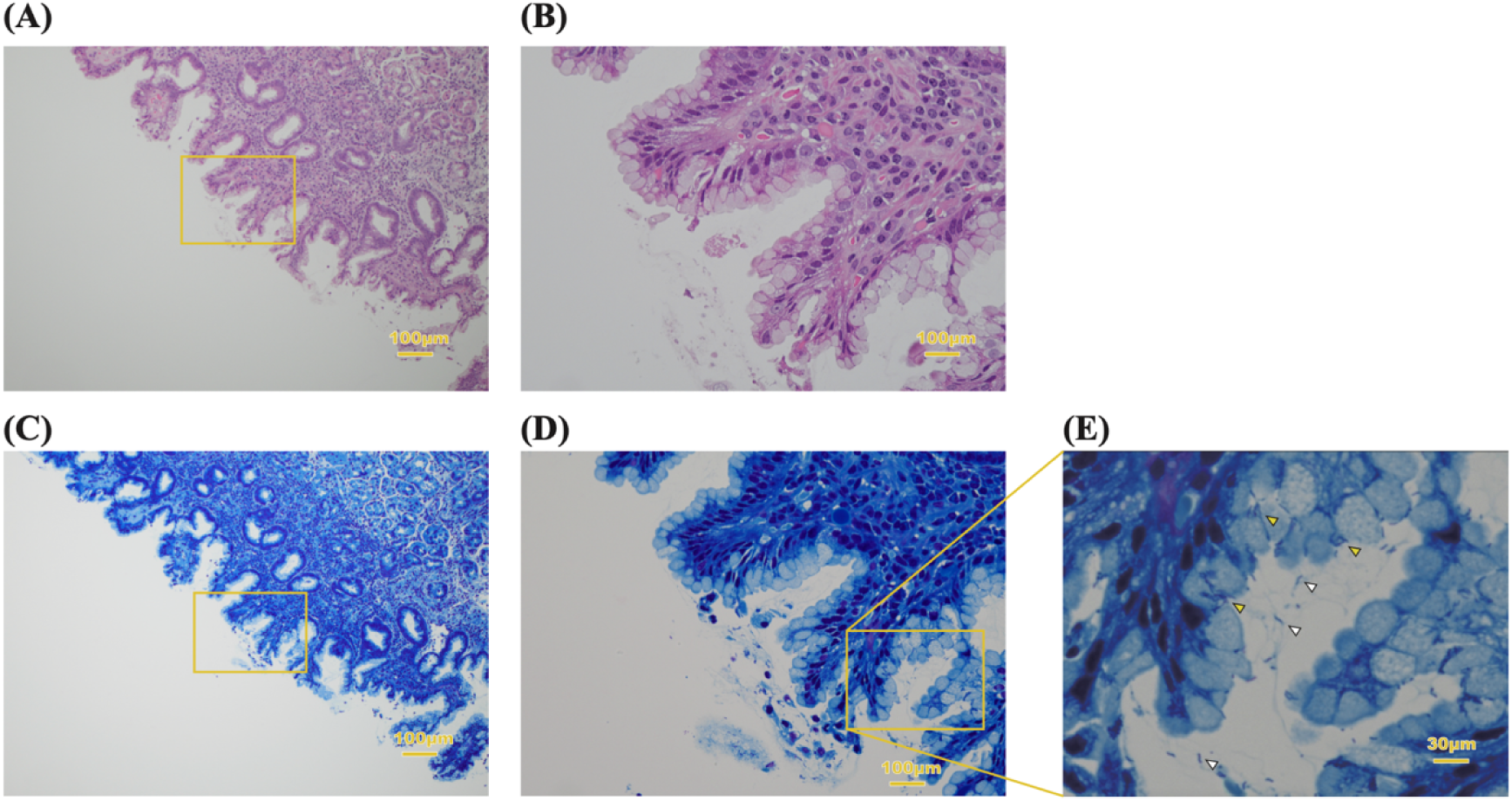
Histopathological observation of biopsy specimen. The gastric mucosa stained with HE (A) and Giemsa (C) of patient no. 15. The original magnification is x 10. (B, D) Higher magnification of the area indicated by the squares in A and C. (E) High-magnification image of the box on the (D). Spiral-like and seagull-like bacilli can be confirmed in the mucus (white arrowhead) and the interior of mucous cells (yellow arrowhead) of the stomach.

### Nested PCR and allele analysis

*H. pylori* DNA was detected in at least one site of the oral cavity in all patients using nested PCR (Table 2). The upper incisors had the highest abundance of the organism (71.4%, p<0.05). Further, *H. pylori* DNA was detected in the biopsy specimens in all patients. And the existence of *H. pylori* DNA was also confirmed in the DNA fragments extracted from an isolate on the agar plate.

**Table 2.** Prevalence of *H. pylori* DNA in gastric tissue and by location in the oral cavity using nested PCR.

135 cavity using nested PCR
| Patient No. | Upper incisor | Lower incisor | Upper right molar | Lower left molar | Saliva | Tongue | Gastric tissue biopsy | Gastric tissue culture |
| --- | --- | --- | --- | --- | --- | --- | --- | --- |
| 1 | + | + | - | - | - | - | + | + |
| 2 | + | - | - | - | - | + | + | + |
| 3 | - | - | + | - | - | - | + | + |
| 4 | + | - | - | - | - | - | + | + |
| 5 | + | - | - | - | - | - | + | + |
| 6 | - | - | + | - | - | + | + | + |
| 7 | - | + | - | - | - | + | + | + |
| 8 | + | - | - | - | - | - | + | + |
| 9 | + | - | + | - | - | - | + | + |
| 10 | + | - | - | - | + | - | + | + |
| 11 | - | - | - | - | - | + | + | + |
| 12 | + | - | - | - | - | - | + | + |
| 13 | + | - | - | - | + | - | + | + |
| 14 | + | - | + | - | - | + | + | + |
| 15 | + | + | - | - | + | - | + | + |
| 16 | + | - | - | - | - | + | + | + |
| 17 | + | - | - | + | - | - | + | + |
| 18 | + | - | - | - | - | - | + | + |
| 19 | + | - | + | - | - | - | + | + |
| 20 | - | + | - | - | - | - | + | + |
| 21 | - | - | - | - | + | - | + | + |
| Prevalence (%) | 71.4 | 19.0 | 23.8 | 4.8 | 19.0 | 28.6 | 100 | 100 |
| n/N | 15/21 | 4/21 | 5/21 | 1/21 | 4/21 | 6/21 | 21/21 | 21/21 |

Table 3 compares the *H. pylori* allelic profiles obtained from oral and gastric samples based on seven housekeeping genes. The number of *H. pylori* allelic profiles ranged from zero to seven, since the yield of DNA was small even when the nested PCR was performed. The alleles of seven loci from both collection sites were determined from only one patient (patient no. 3), and two out of seven alleles matched between oral and gastric samples. Moreover, for one sample set (patient no. 4), no amplification was observed for any of the seven housekeeping genes and therefore, no allele analysis was performed for the sample set. For the rest, allele analysis was performed on the obtained sequences to determine the allele number, of which some were novel, and therefore, unclassified.

**Table 3.** Allele analysis of *H. pylori* DNA sequences obtained from oral and gastric spamples targeting seven house-keeping genes.

| Participants<br>No. | Housekeeping gene |  |  |  |  |  |  |  |  |  |  |  |  |  |
| --- | --- | --- | --- | --- | --- | --- | --- | --- | --- | --- | --- | --- | --- | --- |
|  | *atpA* |  | *urel* |  | *efp* |  | *mutY* |  | *ppa* |  | *trpC* |  | *vacA* |  |
|  | Oral | Gastric | Oral | Gastric | Oral | Gastric | Oral | Gastric | Oral | Gastric | Oral | Gastric | Oral | Gastric |
| 1 | - | - | n/m | 3242 | - | - | - | - | - | - | - | - | - | - |
| 2 | - | - | n/m | 463 | 453 | n/m | - | - | - | - | - | - | - | - |
| 3 | 2364 | 963 | 2643 | 2943 | 1162 | 2421 | 2368 | 2368 | 45 | 942 | 458 | 970 | 573 | 573 |
| 4 | - | - | - | - | - | - | - | - | - | - | - | - | - | - |
| 5 | - | - | - | - | - | - | - | - | n/m | 823 | - | - | - | - |
| 6 | - | - | - | - | - | - | n/m | 2476 | - | - | 970 | 423 | - | - |
| 7 | - | - | n/m | 2647 | - | - | - | - | - | - | - | - | - | - |
| 8 | - | - | n/m | 67 | - | - | - | - | 942 | 929 | - | - | - | - |
| 9 | - | - | 286 | 2856 | - | - | - | - | 849 | 45 | - | - | - | - |
| 10 | 2364 | 1760 | n/m | 466 | - | - | - | - | - | - | - | - | - | - |
| 11 | - | - | - | - | - | - | n/m | 2360 | 420 | 945 | - | - | - | - |
| 12 | - | - | - | - | - | - | - | - | 420 | 448 | - | - | - | - |
| 13 | 2003 | 958 | - | - | - | - | n/m | 2729 | - | - | - | - | - | - |
| 14 | - | - | - | - | - | - | 935 | 38 | - | - | - | - | - | - |
| 15 | - | - | - | - | 453 | 1879 | 935 | 1907 | - | - | - | - | 29 | 950 |
| 16 | 1760 | n/m | 3243 | 966 | - | - | - | - | - | - | - | - | - | - |
| 17 | - | - | n/m | 2471 | - | - | 935 | 2450 | - | - | - | - | - | - |
| 18 | - | - | n/m | 466 | - | - | 3162 | 935 | - | - | - | - | - | - |
| 19 | - | - | n/m | 825 | - | - | 935 | 1219 | - | - | - | - | 785 | 618 |
| 20 | - | - | - | - | - | - | - | - | 1887 | 445 | - | - | - | - |
| 21 | - | - | 738 | 971 | - | - | - | - | - | - | - | - | - | - |

Nested PCR using primers targeting the housekeeping genes did not give amplification products with all the primers. Allele analysis was performed on the obtained sequences to determine the allele sequence types. Some of them were new sequence types that were unclassified (no matching (n/m)).

First, we evaluated the validity of the sequence obtained from the sample by creating a phylogenic tree using the allele type sequence here. Figure 2 shows the phylogenetic analysis of the oral and gastric *H. pylori* based on the allelic profiles of the six housekeeping genes. In addition, a total of 159 isolates from Asia, Oceania, Europe, North America, South America, and Africa were including in the phylogentic analysis. In this analysis, Asia was further categorized as East Asia, Southeast Asia, and South Asia. The *H. pylori* allelic profiles from the oral and gastric samples in this study mainly clustered with the Asian isolates. Partial sequences of the six genes per the patient were shown in Supporting information (S1-S6 File).

**Figure 2.**
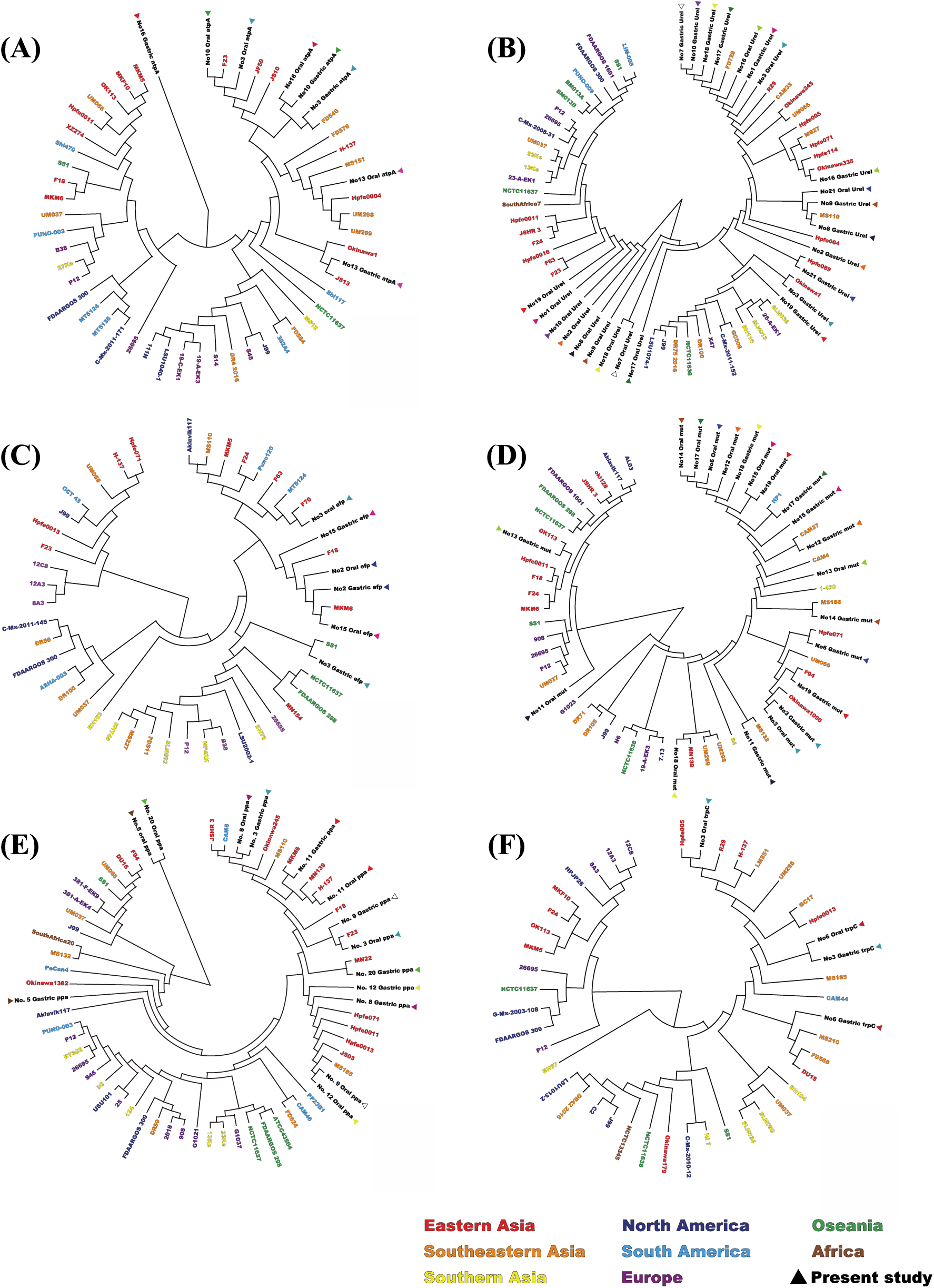
*H. pylori* phylogenetic tree analysis using housekeeping genes. Phylogenetic trees based on the partial sequences of (A) *atpA*, (B) *urel,* (C) *efp*, (D) *mutY*, (E) *ppa*, (F) *trpC*. The 20 strains from this study match the color of the arrowhead so that the oral cavity and stomach correspond to each other. For the 159 strains obtained from BLAST, the color of the letters was changed according to the corresponding region.

### MLST

The genetic relatedness between oral and gastric *H. pylori* was investigated using MLST. DNA sequences of up to six of the housekeeping genes were used for MLST to obtain sequence types. Although the *vacA* gene was able to generate an allele number, it could not be used for MLST analysis on the PubMLST database (http://pubmlst.org/helicobacter/). [29]. Table 4 summarizes the MLST profiles of the sample sets. Of the genotypes obtained from the oral cavity and stomach, only one sample set (patient no. 13) had a matching sequence type. The genotypes of the other 20 sample sets were various and, the gastric and oral *H. pylori* were not identified to be the same.

**Table 4.**
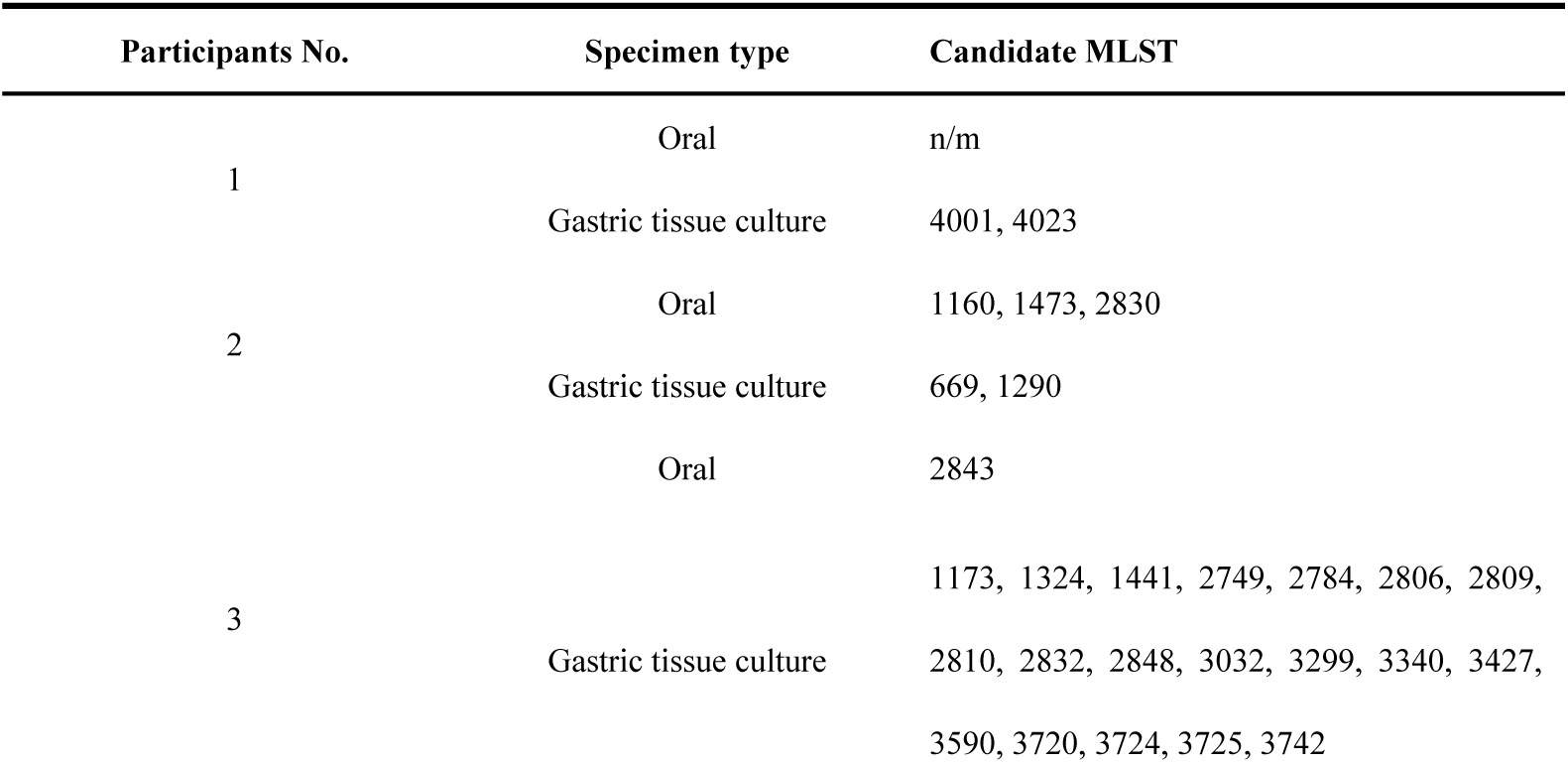
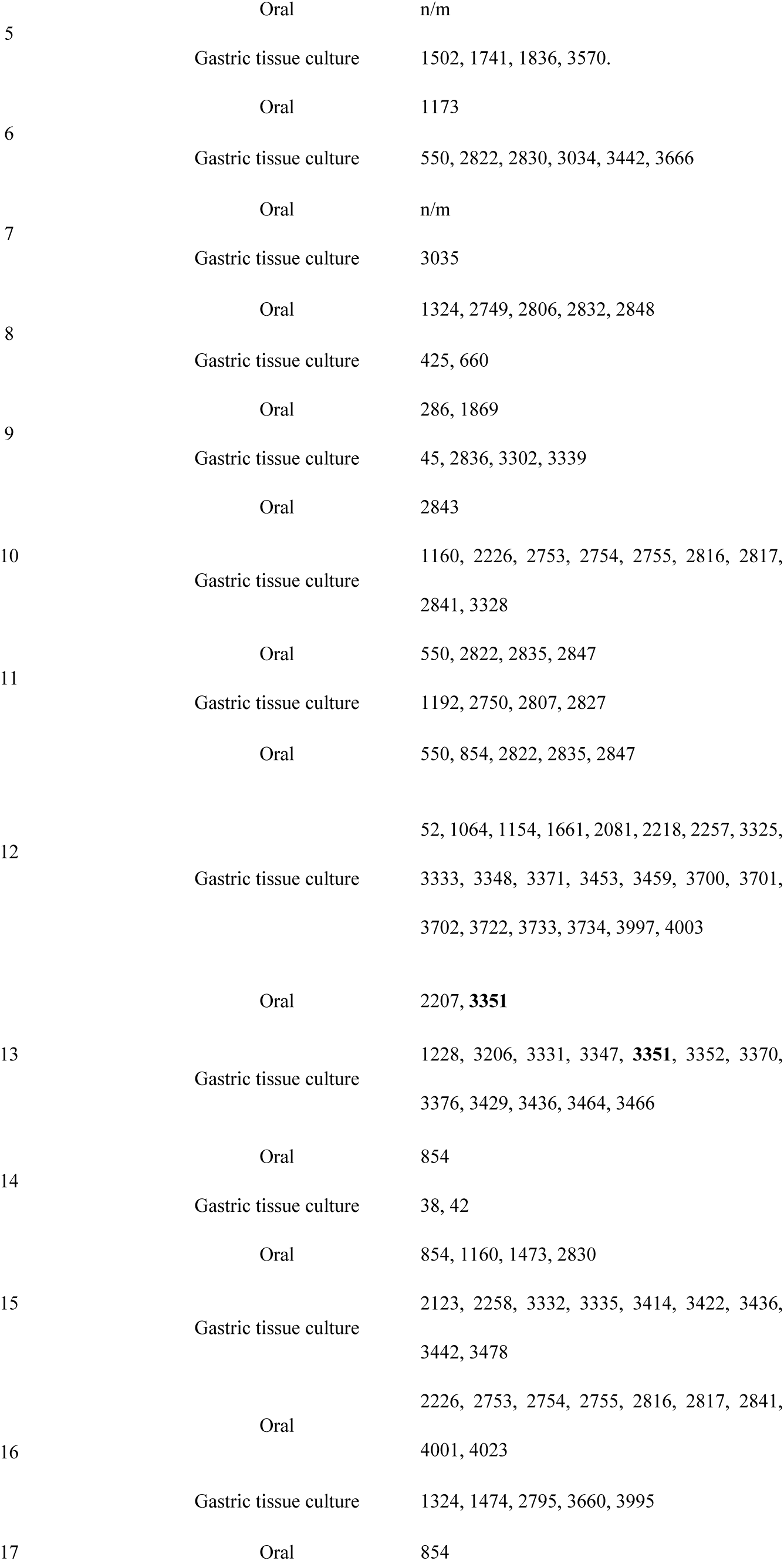
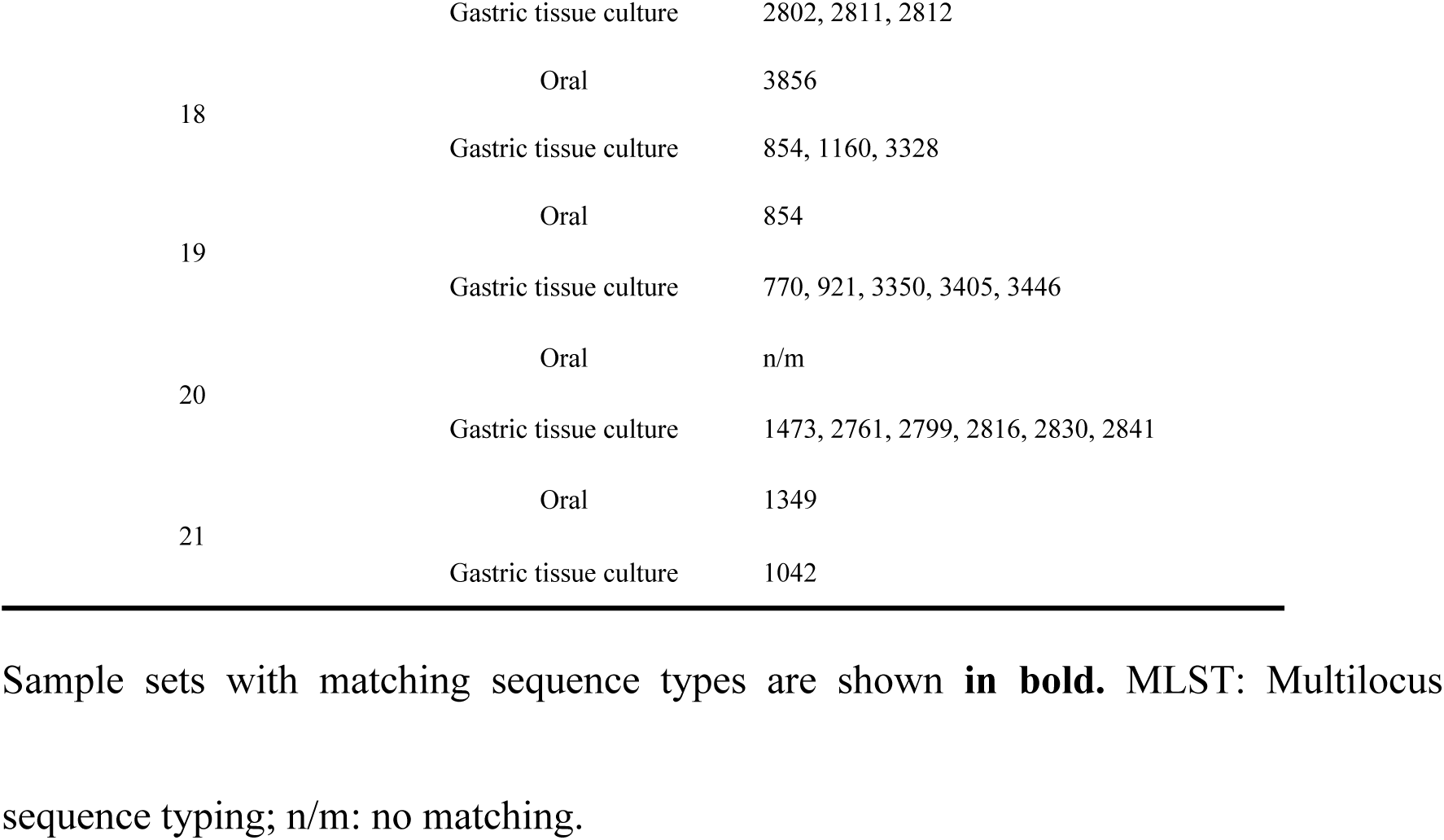
MLST profiles of oral and gastric *H. pylori*.

### Phylogenetic analysis

Phylogenetic tree was created using the samples that obtained a combination of two or more alleles including patient no. 13 (Figure 3). The oral *H. pylori* in patient no. 3 and 15 was markedly similar to gastric *H. pylori*. There was large sequence diversity between two collection sites in patient no. 9, 13 and 19. Partial sequences per the patient were shown in Supporting information (S7 File).

**Figure 3.**
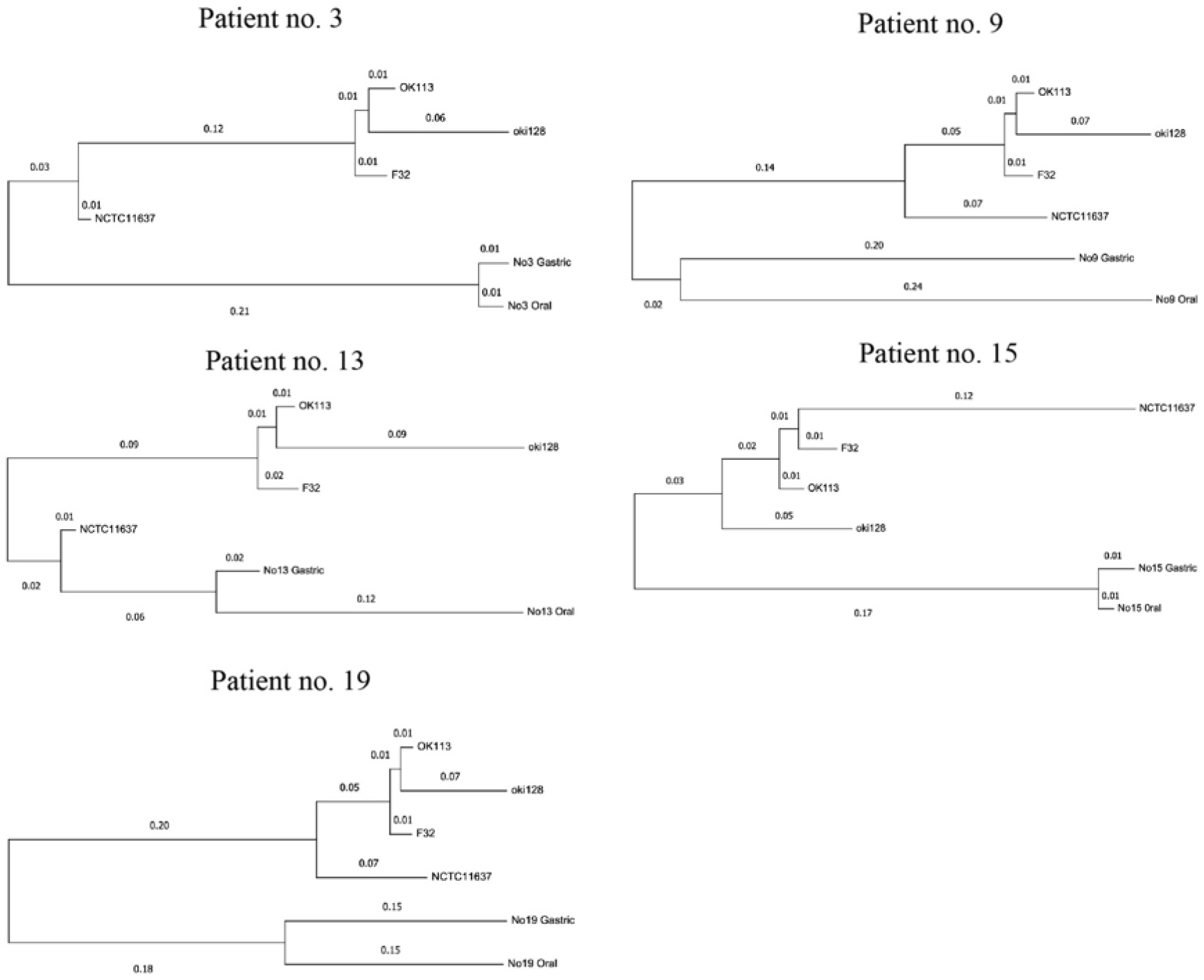
Phylogenetic tree showing the correlation of oral and gastric *H. pylori* based on sequence arrangements. Phylogenetic tree was created using the samples that obtained a combination of two or more alleles including patient no. 13. The gene sequences of *H. pylori* OK113, oki128 and F32, clinical isolates from East Asia, were were downloaded from the BLAST website.

## Discussion

In the present study, we compared the genome sequences of gastric and oral *H. pylori* from 21 patients with early GI cancer using MLST. To our knowledge, this is the first report of the molecular epidemiological analysis of the homology between gastric and oral *H. pylori*. One out of the 21 patients (patient no. 13) harbored the same genotype, indicating that the strains in the stomach and oral cavity may be identical (Table 3). However, the specificity may be low because only two housekeeping genes were used for the analysis. Thus, we created a phylogenetic tree using the samples that obtained a combination of two or more alleles (Figure 3). The result showed that the oral *H. pylori* in two patients was closely related to gastric *H. pylori*, implying the the origins of two strains may be the same. Although there are no reliable reports of successful isolation and culture of oral *H. pylori*, our previous study demonstrated that oral *H. pylori* DNA was detected in the supragingival biofilm collected every two months from one patient who received successful eradication therapy, supporting the existence and persistence of viable *H. pylori* in the oral cavity [12].

In almost all of the phylogenetic trees obtained using the six housekeeping genes, the *H. pylori* sequences in this study clustered with the isolates from the East Asian region (Figure 2). Some *H. pylori* strains have genes involved in carcinogenesis, such as the virulence factors, cytotoxin-associated gene (*cagA*) and vacuumating cytotoxin A (*vacA*) [30, 31]. East Asia, in particular, has a high prevalence of *H. pylori* strains carrying the *cagA* and *vacA* genes, and these are typically virulent strains. Conversely, bacteria with weak expression of these genes are classified as attenuated or non-toxic strains [32]. Therefore, we speculate that both gastric and oral *H. pylori* detected in this study are virulent.

It remains unclear whether *H. pylori* in the oral cavity is an important reservoir for gastric *H. pylori* infections. Most studies that analyzed the homology of *H. pylori* in stomach and oral cavity, compared the genotype of a limited number of pathogenic genes [32–34]. Wang et al. compared *cagA* and *vacA* genotypes of *H. pylori* strains from both saliva and stomach in 31 patients with gastritis and peptic ulcer by PCR. The gastric sample was collected via biopsy from the antrum. The results showed 95% agreement between stomach *H. pylori* isolates and their corresponding saliva DNA in at least one cytotoxin genotype. The authors concluded that the same *H. pylori* strain may exist in the saliva and stomach in the same patient. However, the concordance rate of all four cytotoxin genotypes was only 27%, indicating considerable diversity between two comparison targets. Although DNA sequencing from three patients showed 66.9% to 78% homology of *H. pylori* from both sources, data for the other 28 subjects were not provided [35].

Wongphutorn et al. compared the genotype of *H. pylori* from saliva and stool samples using a partial *vacA* gene sequence. For seven out of 12 individuals, saliva and stool sequences fell into different clusters on a phylogenetic tree, indicating intra-host genetic variation of *H. pylori*. Although this study used only one gene for their comparison, nearly half of the pairs had different genotypes [36].

The discrepancy of genotype between gastric and oral *H. pylori* can be explained by several possible events. One is a mutation in the housekeeping genes, resulting in different allelic profiles. Although housekeeping genes are highly conserved [37, 38], studies have reported that some housekeeping genes can mutate in response to changes in a specific environment [39, 40]. Linz et al. reported that the mutation rate during the acute phase of *H. pylori* infection is more than 10 times faster than during the chronic infection phase [41]. Based on this, it is possible that *H. pylori* which originally invaded the oral cavity may have acquired some gene mutations during infection of the stomach hence the discrepancy in the strain types.

Another possibility is multiple infections with different strains. It is likely that the patient with a *H. pylori* infection in the stomach continued to live in an environment with a high incidence of *H. pylori* infections. Another genotype of *H. pylori* may have infected their oral cavity after the initial infection of stomach. There are also reports of mixed infections of multiple types of *H. pylori* in the oral cavity and stomach [42–44]. Palau et al. reported that, based on housekeeping genes, different strains of *H. pylori* were detected from the same site in the stomach of the same patient. This may also occur in the oral cavity [44]. Thus, future study comparing *H. pylori* DNA extracted from samples taken at different locations is expected.

There is an important limitation to be considered in this study. Although the MLST number is assigned by the combination of partial sequences of six housekeeping genes, 7/21 samples sets were subjected to MLST analysis targeting one housekeeping gene. Moreover although MLST was possible with one gene, the number of candidate MLST numbers increased accordingly, and the specificity and reliability were low. *H. pylori* are present in low abundance in the oral cavity, therefore, despite the use of nested PCR, a sufficient amplification product was not be obtained for further analysis.

In conclusion, different genotypes of *H. pylori* exist in the oral cavity independently of *H. pylori* present in the stomach, there are rare cases in which the same *H. pylori* is present in the stomach and the oral cavity. It is necessary to establish a culture method for oral *H. pylori* for elucidating whether the oral cavity will act as the source of the gastric infection, as our analysis was based on a limited number of combinations of allele sequences.

## Materials and methods

### Participants

The study was conducted at Niigata University Medical and Dental Hospital (Niigata, Japan) between November 2019 and October 2020. We recruited patients with a confirmed *H.pylori* infection using a fecal antigen test who planned to admit for endoscopic surgery on the upper gastrointestinal cancers. The presence of at least one tooth in the oral cavity was included in the inclusion criteria, and participants using denture were not excluded. The study protocol was approved by the Niigata University Ethics Committee (approval number 2019-0220), and the study was carried out in accordance with the approved guidelines. All participants signed an informed consent form before participating in the study.

### Oral sample collection

An oral examination, followed by collection of saliva and biofilm on teeth and tongue, was performed prior to endoscopic surgery. Smoking and oral hygiene status, such as frequency of oral cleaning, and use of oral cleaning aids, were recorded. The number of the DMFT was also recorded. Unstimulated saliva (2 mL) was collected by spitting into a tube. Supragingival dental biofilm samples were collected by scraping from the upper incisors, lower incisors, upper right molars, and lower left molars using a sterile curette. If the participant had no teeth in the designated location, the sample was taken from the opposite site or denture. Each sample was transferred into a tube containing phosphate buffered saline (PBS; pH 7). The superficial layers of the tongue were collected using five gentle strokes from the papillae circumvallatae to the anterior part of the tongue dorsum with a tongue brush (Tongue Cleaner Plus, Ci Medical, Ishikawa, Japan). Bacterial cells were retrieved by vigorous stirring in 20 mL PBS. The samples were centrifuged for 10 min at 10,000 rpm, washed twice with PBS, and stored at −80 °C until further use.

### Gastrointestinal endoscopy and histologic examination

Biopsy specimens were taken from three locations in the stomach using an endoscope: one from the vestibular region and the other two from the body of the stomach. A part of the samples was fixed with 10% formaldehyde and histologically examined by both Giemsa and H-E staining. The pieces were mixed and crushed with a Power masher II (Nippi, Incorporated, Tokyo, Japan) immediately after collection, and half of the amount was inoculated onto *Helicobacter* Selective Agar medium (Nissui Pharmaceutical co., LTD, Tokyo, Japan) and incubated at 37 ℃ under slightly aerobic conditions (5% O2, 10% CO2, and 85% N2), for 72 h. The other half was stored at −80 ℃ until further use.

### DNA extraction and nested PCR

DNA was extracted from the oral samples and biopsy specimens in the stomach as well as from a control strain, *H. pylori* NCTC11637, using the NucleoSpin® Microbial DNA Kit (TaKaRa Bio, Shiga, Japan) according to the manufacturer’s instructions. The quantity and purity of the DNA were assessed by spectrophotometry at 260/280 nm. The DNA was stored at −80 °C until processing.

An aliquot of the DNA extract was amplified using nested PCR as previously described [12], and the presence of *H. pylori* DNA in the sample was confirmed by 1.5% agarose gel electrophoresis. A sample, in which the presence of *H. pylori* DNA was confirmed, was used for the subsequent analysis.

### Sequencing and MLST analysis

MLST was performed as described on PubMLST (http://pubmlst.org/helicobacter/). Primer sets targeting six housekeeping genes, *atpA*, *ureI*, *efp*, *mutY*, *ppa*, and *trpC*, were used (Table 5) [26]. Gene fragments containing these genes were amplified from *H. pylori*-positive specimens by nested PCR.

**Table 5.**
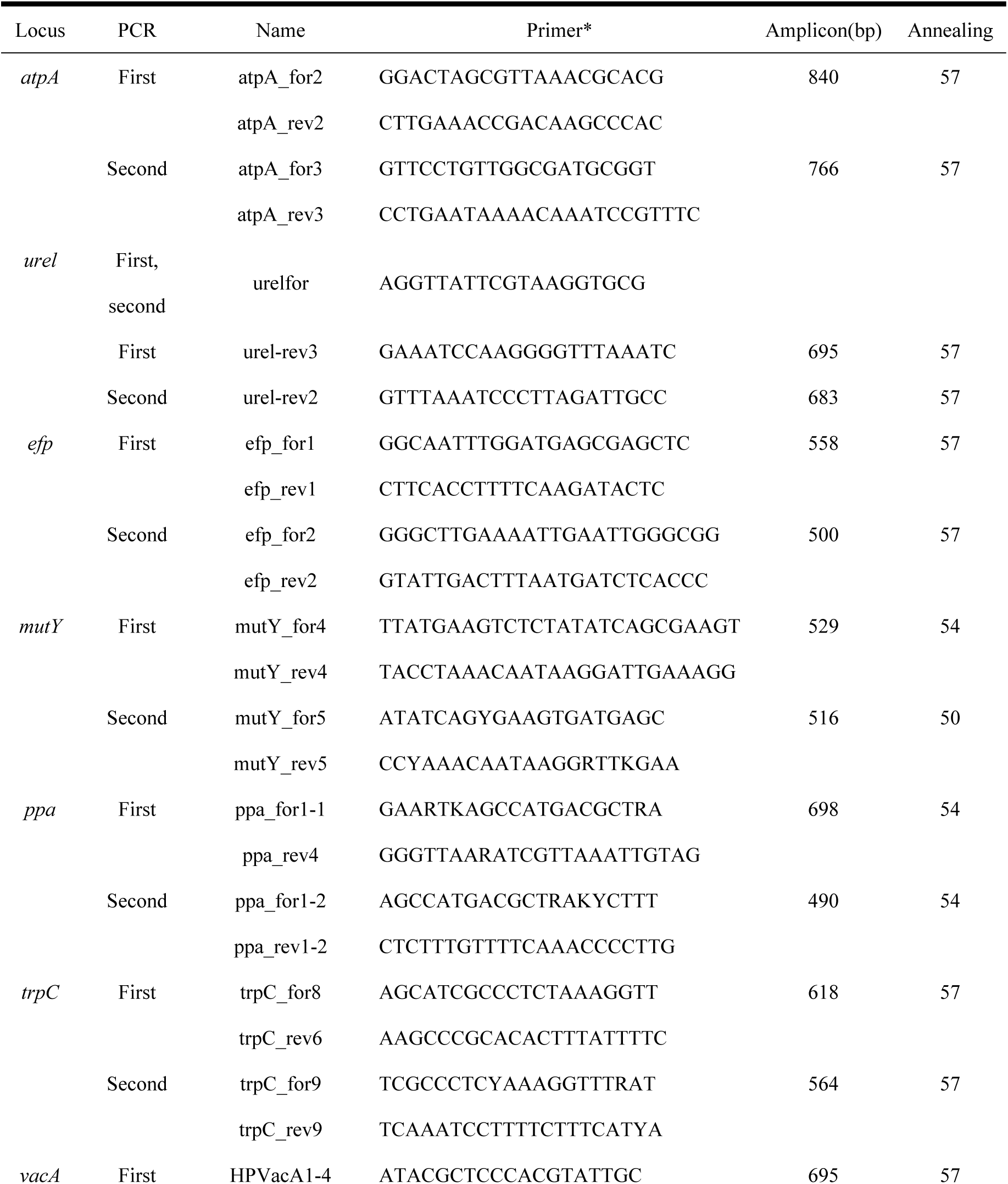
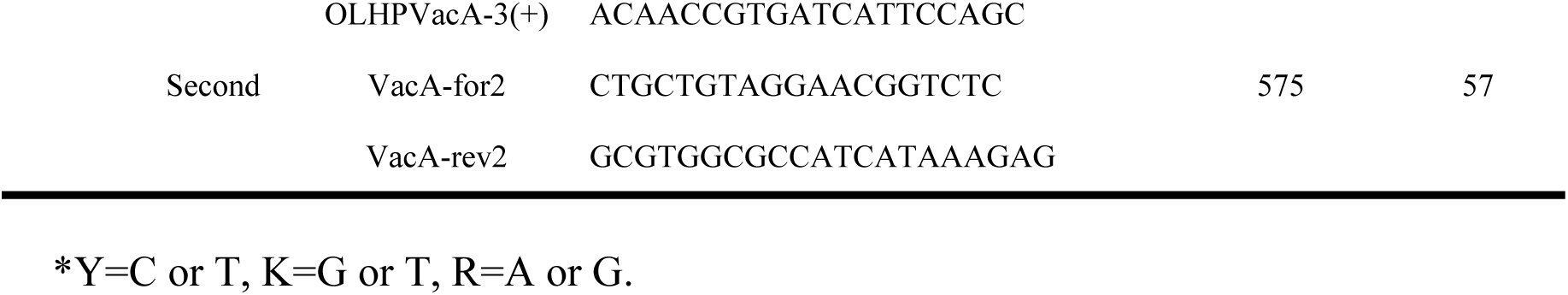
Primers for amplification of housekeeping genes using nested PCR.

Briefly, the first round of PCR amplification was performed using Takara Ex Taq® Hot-Start Version (RR006A; TaKaRa Bio, Shiga, Japan). The amplification comprised 40 cycles of denaturation at 98 °C for 10 s, annealing temperature of each primer for 30 s, and extension at 72 °C for 1 min on a MiniAmp™ Thermal Cycler (Thermo Fisher Scientific, Waltham, MA, USA). For the nested PCR assay, the amplification product (1 µL) obtained by single-step PCR was re-amplified over 40 cycles under the same conditions as in the first round. *H. pylori* NCTC11637 DNA served as the positive control, and water was used as the negative control. Each PCR product was confirmed by 1.5% agarose gel electrophoresis.

The band visualized under LED light was cut out from the gel using a gel band cutter (FastGene ™ □Agarose Gel Band Cutter, Nippon Genetics co., LTD, Tokyo, Japan). Where multiple bands were present, bands of the same size as the positive control band were collected. Amplicons were purified from the gel slices using the Freeze’N squeeze DNA Gel Extraction Spin Clumns (Bio-Rad, Hercules, CA, USA) and NucleoSpin®□ Gel and PCR Clean-up (TaKaRa Bio, Shiga, Japan). The amplified DNA was then sequenced and analyzed using the Applied Biosystems 3730xl DNA analyzer at Macrogen Japan corp. (Tokyo, Japan).

The sequences were uploaded onto PubMLST (http://pubmlst.org/helicobacter/) to determine the closest allele type for each gene. Using the allelic profile of each gene, the sequence type of the sample was determined.

### Phylogenic tree analysis

To analyze the phylogeny of *H. pylori* obtained from the oral cavity and stomach, housekeeping gene sequences of 159 *H. pylori* isolates from 29 countries, in eight regions of the world, were downloaded from the BLAST website (https://blast.ncbi.nlm.nih.gov/Blast.cgi).

Multiple alignment of the MLST genes was then performed using MEGA (V11). The aligned sequences were used to develop a phylogenetic tree. The amplicons obtained from the mouth and stomach were then sequenced and aligned with isolates from Europe, Africa, Asia, America and Oceania using multiple alignments with the MUSCLE program. Asia was classified into East Asia, Southeast Asia, and South Asia. Thereafter, a phylogenetic tree was constructed using the construct/Test Maximum Likelihood tree with the alignment result in MEGA.

### Statistical analysis

Data analysis was carried out using SPSS® 11.0 (SPSS, Chicago, IL, USA). The chi-squared test and Fisher’s exact probability test were used when applicable, and the results were considered statistically significant when the P-value was <0.05. Prevalence was expressed as a proportion and the crude odds ratio (OR) was used to measure the strength of the association between the variables.

## Acknowledgments

The author is grateful to Dr. Tatsuya Ohsumi for his technical support regarding the nested PCR procedure.

## Supporting information

**S1 File. *atpA* DNA sequence of the data used to generate the dendrogram in Figure 2.**

**S2 File. *urel* DNA sequence of the data used to generate the dendrogram in Figure 2.**

**S3 File. *efp* DNA sequence of the data used to generate the dendrogram in Figure 2.**

**S4 File. *mutY* DNA sequence of the data used to generate the dendrogram in Figure 2.**

**S5 File. *ppa* DNA sequence of the data used to generate the dendrogram in Figure 2.**

**S6 File. *trpC* DNA sequence of the data used to generate the dendrogram in Figure 2.**

**S7 File. DNA sequence of the data used to generate the dendrogram in Figure 3.**

